# *Plasmodium falciparum*-specific IgM B cells dominate in children, expand with malaria and produce parasite inhibitory IgM

**DOI:** 10.1101/2020.04.12.030049

**Authors:** Christine S. Hopp, Ababacar Diouf, Kazutoyo Miura, Kristin Boswell, Padmapriya Sekar, Jeff Skinner, Christopher M. Tipton, Michael Chambers, Sarah Andrews, Joshua Tan, Shanping Li, Safiatou Doumbo, Kassoum Kayentao, Aissata Ongoiba, Boubacar Traore, Silvia Portugal, Carole Long, Richard A. Koup, Eric Long, Adrian B. McDermott, Peter D. Crompton

## Abstract

IgG antibodies are known to play a central role in naturally acquired immunity to blood-stage malaria in humans, but little is known about the IgM response to blood-stage malaria, the mechanisms by which IgM may protect, or the underlying biology of *Plasmodium falciparum* (*Pf*)-specific IgM B cells. In a Mali cohort spanning infants to adults we conducted a longitudinal analysis of B cells specific for the *Pf* blood-stage antigens AMA1 and MSP1, as well as the comparator antigen influenza hemagglutinin (HA). At the uninfected baseline, before the malaria season, *Pf*-specific memory B cells (MBCs) in children are disproportionally IgM^+^ and only gradually shift to IgG^+^ with age, in contrast to HA-specific MBCs that are predominantly IgG^+^ from infancy to adulthood. In response to acute febrile malaria, *Pf-*specific IgM B cells increase in frequency and upregulate activation and co-stimulatory markers. B cell receptor (BCR) analysis showed that *Pf*-specific IgM B cells are somatically hypermutated at levels comparable to HA-specific IgG B cells. Finally, IgM antibodies from the plasma of malaria-exposed individuals were comparable to IgG in inhibiting *Pf* blood-stage growth *in vitro*, and significantly better at enhancing phagocytosis of *Pf* merozoites, suggesting that IgM may protect through both direct neutralization and opsonization. Thus, somatically hypermutated *Pf*-specific IgM MBCs dominate in early life, are activated and expand during acute malaria and are associated with plasma IgM that inhibits parasite growth *in vitro*.

## Introduction

Each year there are approximately 200 million cases of *Plasmodium falciparum* (*Pf*) malaria, resulting in nearly half a million deaths (*1*). Human infection is established when a female Anopheles mosquito inoculates the sporozoite stage of the parasite into the skin. Sporozoites enter dermal capillaries and are carried to the liver, where they infect hepatocytes. Over 7-10 days, the parasites replicate without causing symptoms and differentiate into merozoites. Merozoites then exit the liver into the blood stream and infect and replicate within erythrocytes in a 48 hour cycle that rapidy expands the number of parasites in circulation and causes a potentially life-threatening illness. Naturally acquired IgG antibodies are known to play a central role in immunity to blood-stage malaria (*2*), but protective humoral immunity is only acquired after years of repeated infections, likely due to the allelic and antigenic diversity of *Pf* parasites, as well as the relatively short-lived nature of antibody responses to malaria, particulary in children, leaving them susceptible to repeated bouts of febrile malaria (*3*, *4*).

It is well established that long-lasting protective antibody responses require the generation of antibody-secreting long-lived plasma cells (LLPCs) (*5*) and memory B cells (MBCs), that rapidly differentiate into plasmablasts upon rechallenge (*6*). However, studies of children in endemic areas suggest that the response to malaria is dominated by short-lived plasma cells rather than LLPCs (reviewed in (*3*)) and that *Pf*-specific MBCs are acquired relatively inefficiently (*3*, *7*). MBCs can be either isotype switched or express IgM, and although studies have identified a role for IgM in protection against several pathogens (reviewed in (*8*)), the role of IgM antibodies in protection from blood-stage malaria remains unclear.

Several studies have reported *Pf*-specific IgM responses in malaria-exposed populations and in some cases have shown that these responses associate with protection from clinical malaria (*9*–*15*). For example, the malaria-resistant Fulani in West Africa have a higher breadth and magnitude of *Pf*-specific IgM compared to the more susceptible Dogon, whereas *Pf*-specific IgG did not distinguish the two groups (*16*, *17*). More recent studies showed that *Pf*- and *P. vivax*-specific IgM antibodies can mediate complement fixation (*13*, *15*), and that IgM from malaria-experienced individuals can inhibit *Pf* merozoite invasion *in vitro* in the presence of complement (*15*). Furthermore, merozoite-specific IgM antibodies correlated with protection from malaria in a longitudinal cohort of children (*15*). On the other hand, there is evidence that Fcμ-binding proteins expressed on the surface of both *Plasmodium-*infected erythrocytes and merozoites may facilitate immune evasion of the parasite through diverse mechanisms (*18*).

Little is known about the cell biology underlying *Pf*-specific IgM antibody responses. A study in mice found that *P. chabaudi* infection induces long-lasting, somatically mutated IgM MBCs, which dominated the response to a secondary *P. chabaudi* infection (*19*). The same study showed evidence of *Pf*-specific IgM MBCs in naturally exposed human individuals, yet the dynamics and biology of *Pf*-specific IgM B cells in humans remain unclear. Here, we generated *Pf* antigen B cell probes specific for the apical membrane antigen 1 (*Pf*AMA1) and merozoite surface protein 1 (*Pf*MSP1) and performed a longitudinal analysis of naturally acquired *Pf*-specific B cells in response to acute febrile malaria, as well as a cross-sectional analysis of *Pf*-specific B cell in children and adults. To provide insight into the relative role of *Pf* in driving the observed phenotypes, influenza hemagglutinin (HA)-specific B cells were tracked simultaneously in the same individuals.

We found that *Pf*-specific MBCs in children are disproportionally IgM^+^ and only gradually shift to IgG^+^ with age, in contrast to HA-specific MBCs that are predominatly IgG^+^ from infancy to adulthood. In response to acute febrile malaria *Pf-*specific IgM B cells increase in frequency and upregulate activation and co-stimulatory markers. B cell receptor (BCR) analysis showed that *Pf*-specific IgM B cells are somatically hypermutated at levels comparable to HA-specific IgG B cells. We further show that purified total IgM antibodies from plasma of malaria-exposed individuals inhibit parasite growth *in vitro* in the absence of complement, and that IgM antibodies are superior to IgG antibodies in mediating opsonic phagocytosis of *Pf* merozoites. This analysis provides new insights into the mechanisms by which IgM may protect from malaria as well as the underlying biology of *Pf*-specific IgM B cells during natural infection.

## Results

### Confirmation of specificity of *Pf*AMA1/*Pf*MSP1 and HA B cell probes

*Pf*AMA1 and *Pf*MSP1 are immunogenic proteins (*20*) expressed on the surface of *Pf* blood-stage merozoites and are known to be involved in merozoite attachment and invasion of erythrocytes (reviewed in (*21*). As a comparator non-malaria antigen, we used influenza surface glycoprotein hemagglutinin (HA)—the principal target of influenza-specific neutralizing antibodies. Using the Meso Scale Discovery (MSD) assay, which is an electrochemiluminescent-based ELISA replacement assay (*22*), we found in the Mali cohort a high prevalence of serum IgG reactivity specific for the influenza A subtypes H1N1 or H3N2, which have been reported to circulate in Mali (*23*) (Fig. S1). Of note, the influenza-specific B cells studied here are naturally acquired, since influenza vaccine programs have yet to be widely implemented in Mali (*24*). To track *Pf*- and HA-specific B cells, we generated antigen probes using recombinant *Pf*MSP1 and *Pf*AMA1 (*25*, *26*) and trimeric HA (subtypes H1 and H3; (*27*)), respectively. Biotinylated proteins were coupled to fluorescently labelled streptavidin and the resulting probes were used as previously described (*27*) for *ex vivo* flow cytometry of cryopreserved PBMCs. Each probe was labelled with two colors, such that single color positive cells likely representing binding to the fluorophore were excluded (Fig. 1A-D). PBMCs were obtained from subjects aged 3 months to 36 years residing in the malaria endemic village of Kalifabougou, Mali. Details of the cohort study design and study site have been described in detail previously (*4*). PBMCs were collected at the end of the six-month dry season in May, a timepoint referred to as “healthy baseline” and one week after treatment of the first febrile malaria episode of the ensuing 6-month malaria season, at “convalescence”. PBMCs were stained simultaneously with *Pf*AMA1 and *Pf*MSP1 probes in order to increase the frequency of *Pf*-specific B cells detected in any given sample, thus *Pf*AMA1 and *Pf*MSP1 probe-binding cells are indistinguishable by flow cytometry analysis in this study and are together referred to as “*Pf*^+^“cells.

**Fig. 1.**
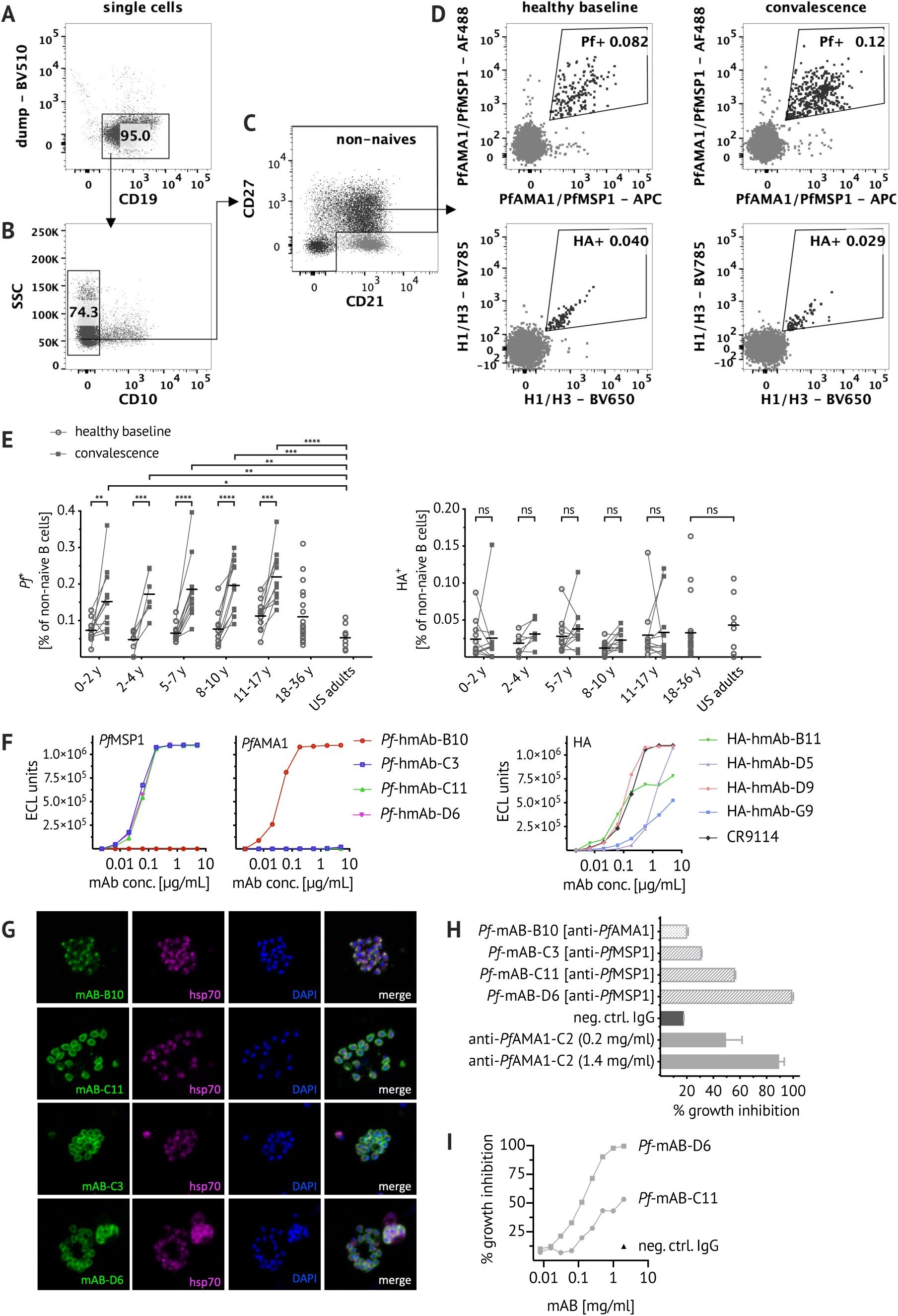
Confirming specificity of *Pf*AMA1/*Pf*MSP1 and HA B cell probes. Representative flow cytometry plots of Malian PBMCs after B cell enrichment showing gating strategy to exclude dump channel positive non-B cells (CD3^+^ CD4^+^ CD8^+^ CD14^+^ CD16^+^ CD56^+^) (**A**), CD10^+^ immature B cells (**B**) and CD21^+^ CD27^−^naïve B cells (**C**). (**D**) *Pf*AMA1 or *Pf*MSP1 (*Pf*^+^) and influenza (HA^+^) probe-binding cells were identified longitudinally at healthy baseline and convalescence. (**E**) Frequencies of *Pf*^+^/HA^+^ cells at healthy baseline (circles) and convalescence (squares) in Malian children aged 0-2 years (n=11), 2-4 years (n=7), 5-7 years (n=11), 8-10 years (n=11), 11-17 years (n=11); as well as healthy Malian (n=20) and U.S. adults (n=8). Each dot indicates an individual, connected lines show paired samples, while bars show means. (**F**) *Pf*^+^ and HA^+^ cells were single-cell sorted from adult Malian donors and their matched heavy and light chain BCR sequences were used to express human monoclonal antibodies (hmAbs) (see suppl. Table 1 for VDJ gene usage). HmAbs were tested for binding to *Pf*MSP1/*Pf*AMA1 and HA using the MSD assay. Both H1 and H3 HA were used but only binding to H1 HA was detected and is displayed here. Binding of four *Pf*-hmAbs (*Pf*-B10, *Pf*-C3, *Pf*-C11 and *Pf*-D6), four HA-hmAbs (HA-B11, HA-D5, HA-D9, HA-G9) and the HA-specific positive control hmAb CR9114 (*29*) is shown. (**G**) For immunofluorescent analysis, *Pf-*3D7 schizonts were stained with *Pf*-hmAbs (green, first column), co-stained with anti-hsp70 antibody (purple, second column) and the nuclear stain DAPI (blue, third column). (**H**) Biological activity of *Pf*-hmAbs was assessed via *in vitro* GIA at 2 mg/ml and compared to the positive control polyclonal rabbit anti-*Pf*AMA1-C2 (*31*). (**I**) Growth inhibitory activity of *Pf*-hmAbs C11 and D6 was tested at concentrations ranging from 0.0078 to 2 mg/ml. Statistical analysis: (E) 2way ANOVA with Bonferroni’s multiple comparisons test.

After exclusion of non-B cells (CD3^+^ CD4^+^ CD8^+^ CD14^+^ CD16^+^ CD56^+^), immature (CD10^+^) and naïve (CD21^+^ CD27^−^) B cells (Fig. 1A-C), the frequency of *Pf*^+^ cells (Fig. 1D) ranged from an average of ~0.07 % of total non-naïve B cells in children under 5-year of age to ~0.1 % in adults at the healthy basline (Fig. 1E), although the trend toward increased numbers of *Pf*^+^ cells with increasing age was not statistically significant. Of note, the total population of non-naïve B cells was stable across age groups and from healthy baseline to convalescence. Across all age groups, *Pf*^+^ cells increased by approximately two-fold at the convalescent time point (Fig. 1E). The frequency of HA^+^ B cells was lower, comprising approximately 0.025% of total non-naive B cells in both Malian and U.S. adults (Fig. 1E). No heterologous boosting of HA^+^ B cells at convalescence was observed (Fig. 1E).

To more directly confirm the antigen specificity of the B cell probes, we sorted single probe-positive B cells from adult Malian PBMC samples (gating strategy in Fig. 1 A-D) and amplified the BCR using reverse transcription and PCR followed by Sanger sequencing (*22*, *28*). Heavy chain and corresponding light chain sequences of four *Pf*- and HA-probe binding B cells were chosen for cloning and expression as recombinant human monoclonal antibodies (hmAb). Information on VDJ gene usage, CDR3 sequence and the number of somatic mutations of cloned hmAbs is shown in supplementary Table 1. Binding of these hmABs was characterized using the MSD assay: three of the *Pf*-hmAbs recognized *Pf*MSP1, while one (clone B10) bound to *Pf*AMA1 (Fig. 1F). HA-specific hmAbs did not bind to H3 HA, but bound to H1 HA with variable affinity, with one of the HA-hmAbs (clone B11) displaying a binding curve comparable to the HA-specific positive control hmAbs CR9114 (Fig. 1F) (*29*). The specificity of *Pf*-hmAbs was further confirmed by immunofluorescent assays (IFA) using *Pf* blood stage parasites (3D7) grown *in vitro* (Fig. 1G). The *Pf*-hmAb clone B10, which bound to *Pf*AMA1 in MSD assays, had a micronemal staining pattern, consistent with the cellular location of *Pf*AMA1, while the three *Pf*MSP1-specific hmAbs stained the merozoite surface as expected (Fig. 1G). The functional activity of *Pf*-hmAbs was evaluated by a standardized *in vitro* parasite growth inhibition assay (GIA) (Fig. 1 H-I), as described previously (*30*). The *Pf*MSP1-specific hmAbs, that exhibited similar binding curves in the MSD assay had varying levels of parasite growth inhibitory activity, with clone D6 inhibiting as well as the polyclonal anti-*Pf*AMA1 positive control (Fig. 1H). The *Pf*AMA1-specific clone B10 had no inhibitory activity.

### *Pf*-specific MBCs in children are skewed toward IgM

The validated *Pf*- and HA-B cell probes were then used to analyze the isotype of antigen-specific B cells by flow cytometry (gating strategy in Fig. 2A-G). After exclusion of naïve (CD21^+^ CD27^−^) B cells (Fig. 2A) an average of 55% of *Pf*^+^ B cells were IgM in children aged 5-17 years (n=33), while an average of only 20% of HA^+^ B cells were IgM in the same subjects (Fig. 2H). Of non-naïve *Pf*^+^ IgM^+^ B cells, just over 20% were IgD-(Fig. 2I) and just over 40% expressed IgD and CD27 (Fig. 2J). Even after exclusion of IgD^+^ B cells, a substantial percentage of *Pf*^+^ IgD-B cells were not isotype switched—40% in 0-2-year olds and 20% in children older than 2 years (Fig. 2K). This was in contrast to IgD-HA^+^ B cells which were primarily IgG across all age groups (Fig. 2L). A possible explanation for the high prevalence of unswitched B cells could be asymptomatic parasitemia, as chronic persistence of blood stage parasites might drive continuous B cell stimulation. Indeed, in Kalifabougou the prevalence of asymptomatic parasitemia is approximately 50% at the end of dry season (*32*); however, we found that there is no difference in the isotype distribution of *Pf*-specific B cells at the end of the dry season in uninfected and asymptomatically infected individuals (Fig. 2M). Of note, switched and unswitched *Pf*^+^ IgD-B cells had comparable distributions of classical MBCs (CD21^+^CD27^+^), atypical MBCs (CD21^−^CD27^−^) and activated MBCs (CD21^−^CD27^+^) (Fig. S2A). Given that over half of circulating non-naïve *Pf*^+^ B cells were unswitched, we further investigated the phenotype and function of *Pf*^+^ IgM^+^ B cells.

**Fig. 2.**
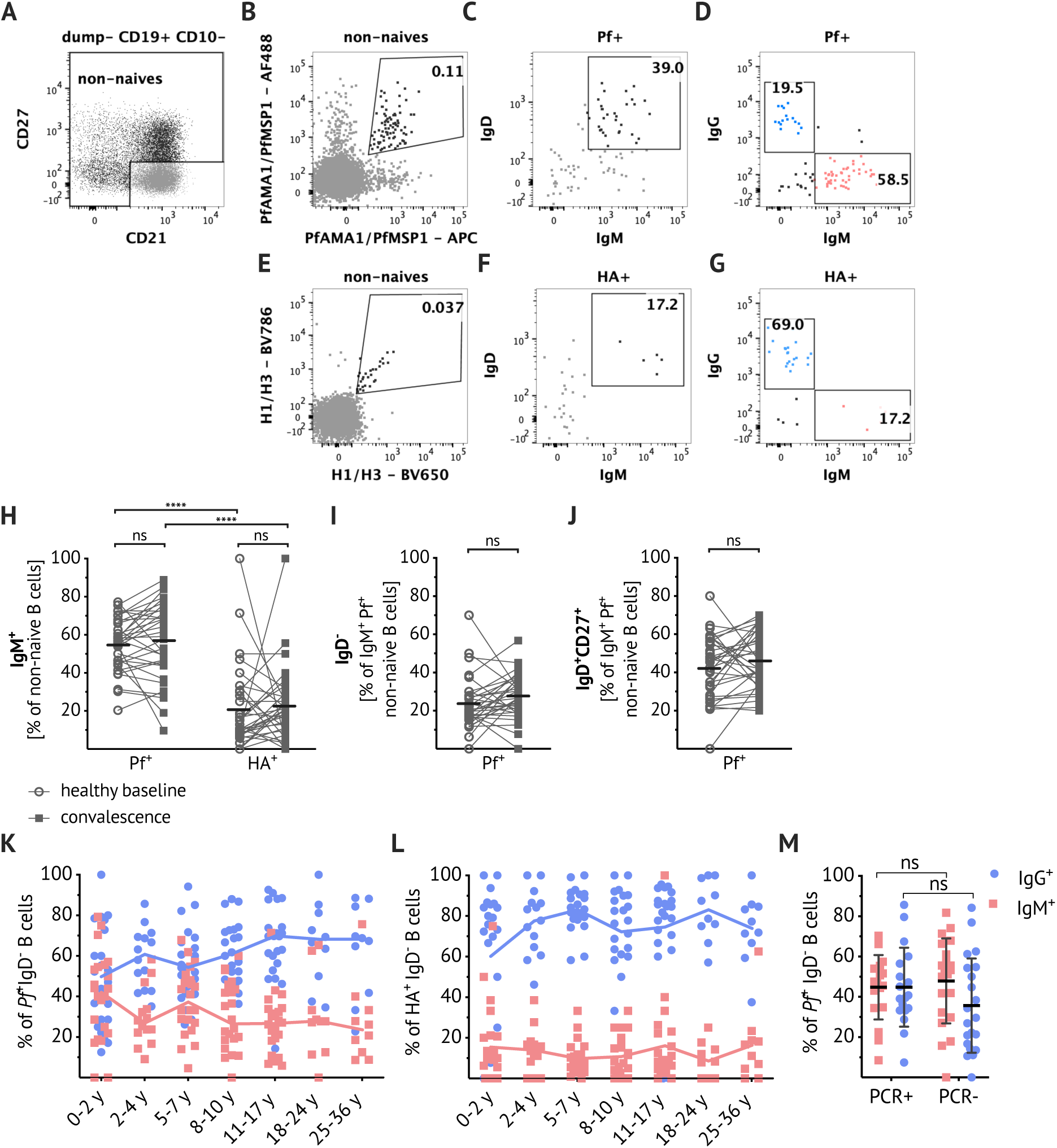
*Pf*-specific MBCs are skewed toward IgM. (**A-G**) Representative plots showing dump channel-negative (CD3^+^ CD4^+^ CD8^+^ CD14^+^ CD16^+^ CD56^+^) CD19^+^ CD10-B cells identified in PBMCs of Malian subjects. After exclusion of naïve (CD27^−^CD21^−^) B cells (**A**), gating on *Pf*^+^ (**B**) and HA^+^ (**E**) probe-positive B cells and IgD/IgM staining (**C&F**), as well as IgG/IgM staining (**D&G**) within the respective antigen-specific B cells is shown. (**H**) Percentage of IgM^+^ B cells within *Pf*^+^ and HA^+^-specific B cells at healthy baseline (circles) and convalescence (filled squares). Percentages of IgD-(**I**) and IgD^+^ CD27^+^ (**J**) within IgM^+^ *Pf*^+^-specific B cells at healthy baseline (circles) and convalescence (filled squares). (**H-J**) Connected lines show paired samples, while bars show means. Data obtained from 5-17 year old Malians (n=33; average age of the subjects was 9.6 years and 42% were female). (**K&L**) Percentage of IgM^+^ (red squares) and IgG^+^ (blue circles) B cells within IgD-Pf^+^ (**K**) or IgD-HA^+^ (**L**) populations; combined data from healthy baseline and convalescence are shown for subjects aged 0-2 years (n=11), 2-4 years (n=7), 5-7 years (n=11), 8-10 years (n=11), 11-17 years (n=11); and data from healthy baseline shown for subjects aged 18-24 years (n=10) and 25-36 years (n=10). Line connects median. (**M**) Percentage of IgM^+^ (red squares) and IgG^+^ (blue circles) B cells within the IgD^-^Pf^+^ population, in individuals with available *Pf* PCR data: healthy 13-15-year-olds were stratified by the presence (PCR^+^, n=18) or absence (PCR-, n=21) of asymptomatic *Pf* parasitemia at the end of the dry season. Statistical analysis: (H&M) 2way ANOVA with Bonferroni’s multiple comparisons, (I-J) Wilcoxon matched pairs signed rank test.

### Somatic hypermutation rates of *Pf*-specific IgM B cells and HA-specific IgG B cells are similar

Somatic hypermutation and affinity maturation are key mechanism in the generation of antibody diversity whereby point mutations accumulate in the variable regions of the heavy and light chain of immunoglobulin genes and B cells with high-affinity for their cognate antigen are positively selected. To gain insight into the developmental history of *Pf*^+^ IgM B cells and their relationship to isotype switched *Pf*^+^ B cells, the *Pf*-specific BCR repertoire was interrogated. *Pf*^+^ and HA^+^ B cells were single-cell sorted from PBMCs of healthy Malian adults with a lifelong exposure to intense *Pf* transmission, as well as healthy malaria-naïve U.S. adults (see Fig. S3A for sorting strategy). Heavy chain gene sequences were obtained from 328 *Pf*-specific and 165 HA-specific B cells, and corresponding light chain gene sequences were obtained from 139 *Pf*-specific and 76 HA-specific B cells. The numbers of somatic mutations in both heavy and light chain sequences were calculated with IMGT/V-QUEST (*33*), using germline BCR sequences available on IMGT as the reference. *Pf*^+^ IgG BCRs had an average of 32 mutations per heavy chain while HA^+^ IgG BCRs had 19 mutations per heavy chain on average (Fig. 3A). On average *Pf*^+^ IgM BCRs had fewer mutations than their *Pf*^+^ IgG counterparts, but with a mean of 16 mutations per *Pf*^+^ IgM heavy chain, they were comparable to HA^+^ IgG BCRs (Fig. 3A). Of note, *Pf*^+^ IgM BCRs isolated from Malian individuals had higher numbers of mutations than those obtained from healthy malaria-naïve U.S. adults (mean of 9 mutations per heavy chain), indicating that *Pf*^+^ IgM BCRs in healthy malaria-exposed adults are MBCs. The distribution of mutations across the cell subsets was similar for heavy and light chains, although the overall mutation load was lower for light chains (Fig. 3B). There were insufficient *Pf*^+^ IgG B cells isolated from malaria-naïve U.S. adults for analysis. The *Pf*^+^ IgM B cells with the highest BCR mutation rates were among CD21^−^CD27^−^ atypical MBCs (Fig. S2C), a B cell subset that expands in association with malaria transmission (*34*, *35*). V and J gene usage of Malian *Pf*^+^ IgM and IgG BCRs were analyzed and while IgHV3-30 and IgHV4-59 were frequent, a wide range of V or J genes were present, indicating that a large number of V and J genes can bind *Pf*AMA1 or *Pf*MSP1 (Fig. 3C).

**Fig. 3.**
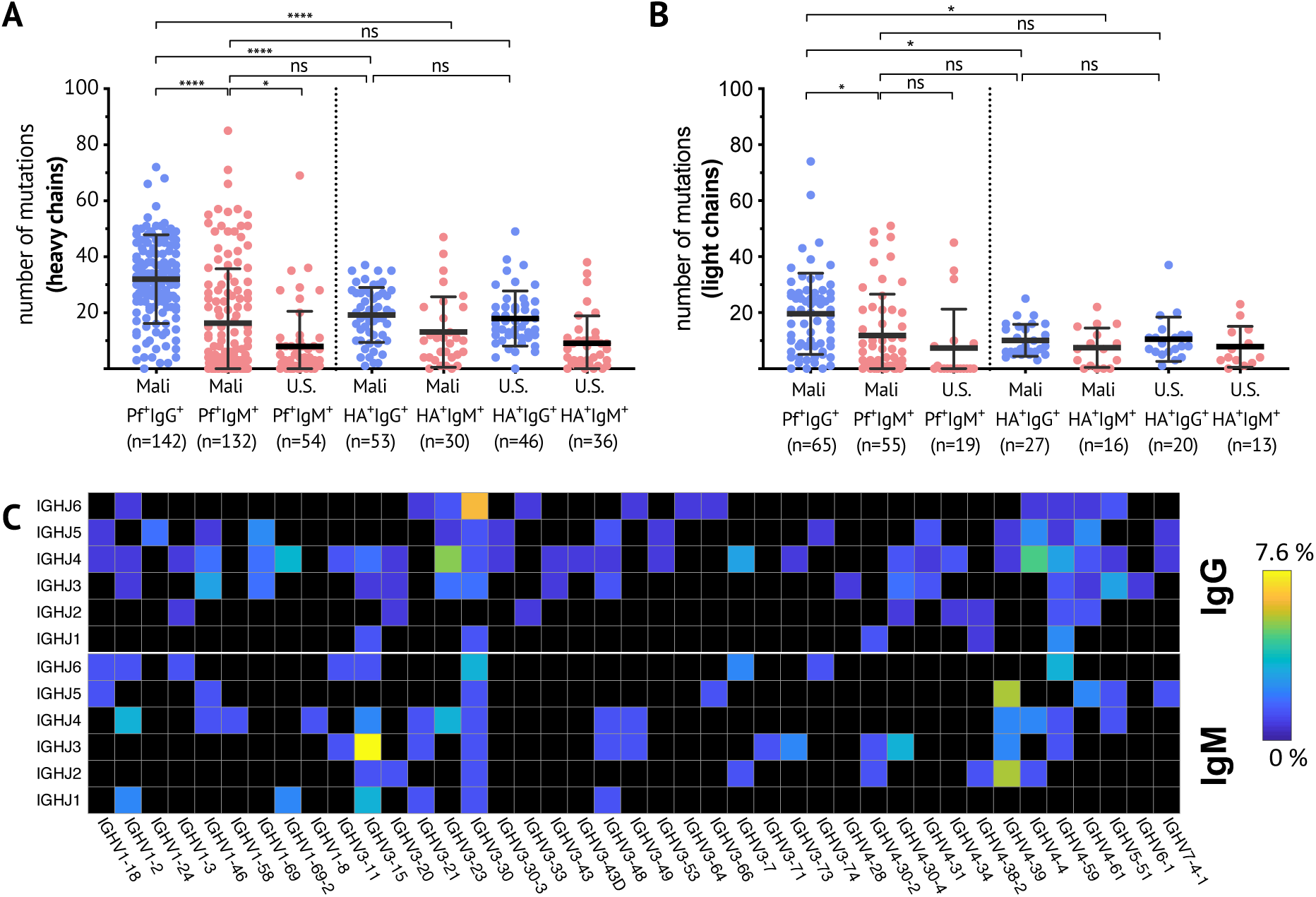
Somatic hypermutation analysis of *Pf*^+^ and HA^+^ B cells. Number of mutations in the variable region of the heavy chain (**A**) and light chain (**B**) of individual *Pf*^+^ or HA^+^ IgG (blue) or IgM (red) B cells. Data combined from five healthy adult Malian donors and three healthy U.S. donors. Each dot indicates a single cell; lines and whiskers represent means and SDs, respectively. (**C**) Heat map showing usage of Ig heavy chain variable (IGHV) and joining (IGHJ) genes, displayed as percentage of total sequences, of *Pf*^+^ IgG (top panel) and *Pf*^+^ IgM (lower panel) BCRs. Statistical analysis: (A-B) One-way ANOVA with Tukey’s multiple comparisons test.

### *Pf*-specific IgM B cells are activated and expand during acute febrile malaria

To understand whether and how *Pf*-specific IgM B cells participate in the immune response to acute *Pf* infection, *Pf*^+^ IgM B cells were profiled *ex vivo* by flow cytometry for expression of markers of B cell activation and surface markers associated with T cell interaction. The frequency of activated (CD21^−^CD27^+^) *Pf*^+^ IgM^+^ B cells increased seven days after treatment of acute febrile malaria (convalescence), relative to the healthy baseline before the malaria season (Fig. 4A, G). Furthermore, *Pf*^+^ IgM B cells upregulated CD86 (B7.2) at convalescence relative to baseline (Fig. 4B, I). CD86 is a marker of activated B cells (*36*) and a costimulatory ligand for CD28, suggesting that activated *Pf*^+^ IgM B cells may interact with T cells. At convalescence, MHC class II was more clearly upregulated on *Pf*^+^ IgG B cells compared to *Pf*^+^ IgM B cells (Fig. 4C, K). CD71 (transferrin receptor 1), a membrane glycoprotein that plays an important role in cellular uptake of iron, is upregulated in cells that require additional iron for cell differentiation and proliferation (*37*). A higher frequency of CD71^+^ *Pf*^+^ IgM B cells was observed at convalescence (Fig. 4D, M), suggesting metabolic activation of *Pf*^+^ IgM B cells. Furthermore, the proliferation/activation marker Ki67 increased in *Pf*^+^ IgM B cells at convalescence (Fig. 4E, O). A significant increase in the frequency of *Pf*^+^ IgM B cells expressing the lineage-defining transcription factor T-bet was seen at convalesence (Fig. 4F, Q). High levels of T-bet have previously been correlated with IgG3 class switching in atypical MBCs in malaria-exposed individuals (*38*), suggesting that T-bet^+^ *Pf*^+^ IgM B cell may be primed to switch to IgG3. Importantly, while CD21^−^CD27^+^, Ki67^+^ and T-bethi *Pf*^+^ IgM B cells increased significantly at convalescence, the frequency of corresponding HA^+^ IgM B cell subsets remained unaffected, indicating a relative lack of heterologous activation of HA^+^ IgM B cells in response to febrile malaria. Although there was an increase in CD86^+^ and CD71^+^ HA^+^ IgM B cells at convalescence, it is important to note that the frequency of HA^+^ IgM B cells is lower than *Pf*^+^ IgM B cells (Fig 2H).

**Fig. 4.**
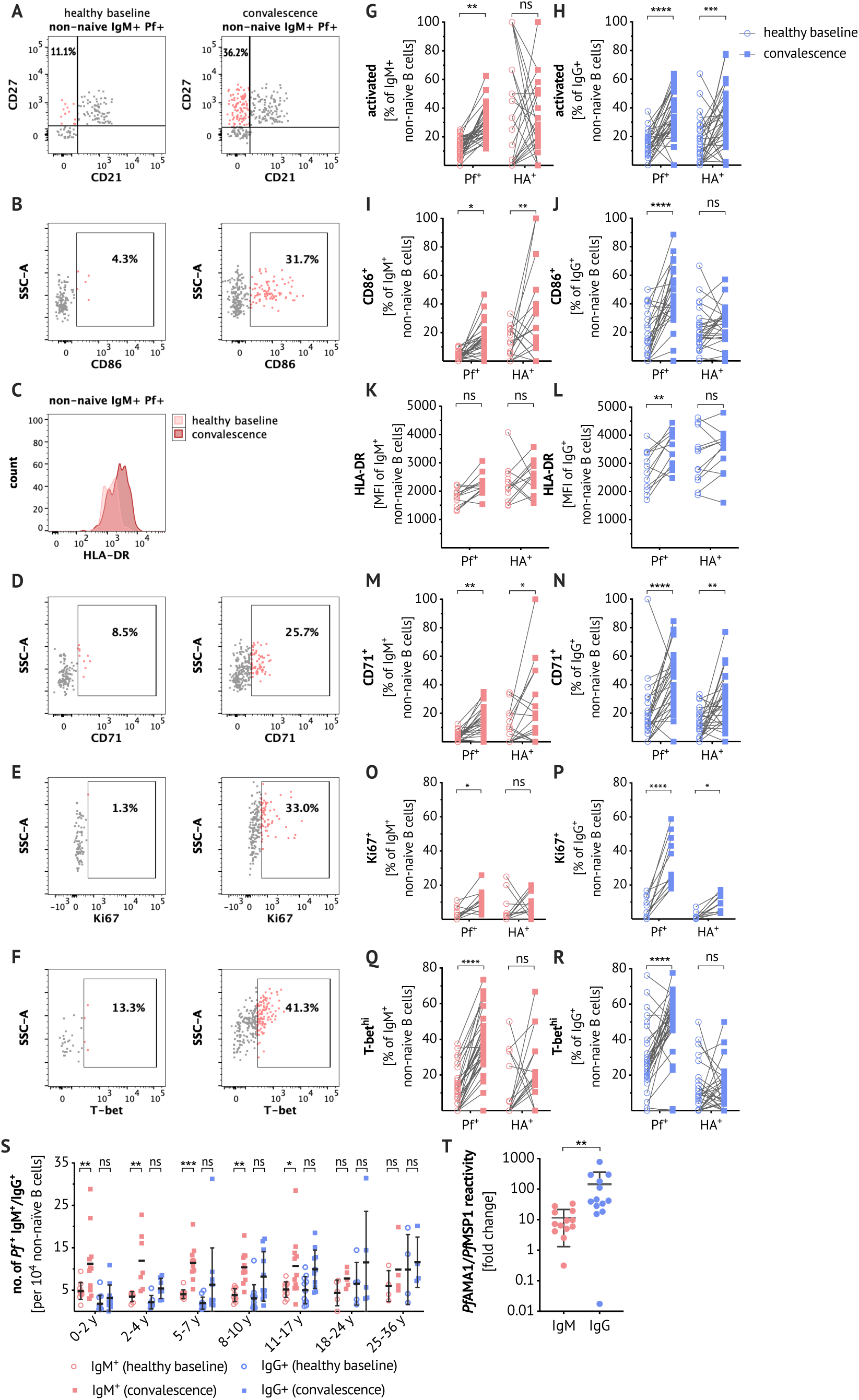
*Pf*-specific IgM B cells are activated, proliferate and expand in response to febrile malaria. Representative plots showing gating on activated (CD27^+^CD21^−^) B cells (**A**), CD86^+^ (**B**), CD71^+^ (**D**), Ki67^+^ (**E**) and T-bethi (**F**) B cells and mean fluorescence intensity (MFI) of HLA-DR staining (**C**) within the *Pf* ^+^ IgM^+^ B cell population at baseline and convalescence after excluding CD27^−^CD21^+^ (naïve), CD10^+^ immature B cells and CD3^+^ CD4^+^ CD8^+^ CD14^+^ CD16^+^ CD56^+^ non-B cells. Panels (**G-S**) show *ex vivo* frequencies and expression levels in antigen-specific B cell populations only. Frequency of activated (**G-H**) or CD86^+^ (**I-J**), MFI of HLA-DR staining (**K-L**), frequency of CD71^+^ (**M-N**), Ki67^+^ (**O-P**) and T-bethi (**Q-R**) in *Pf*^+^ or HA^+^, IgM^+^ or IgG^+^ B cells *ex vivo* at healthy baseline (circles) and convalescence (squares) are shown. Data from 5-17 year-olds is merged to reduce figure complexity; (n=33) (G-H & Q-R), (n=12) (K-L & O-P), (n=27) (I-J & M-N). (**S**) Number of *Pf*^+^ IgM (red) or *Pf*^+^ IgG (blue) B cells per 104 CD10-non-naïve B cells at baseline (circles) and convalescence (squares) by age: 0-2 years (n=11), 2-4 years (n=7), 5-7 years (n=11), 8-10 years (n=11), 11-17 years (n=11), 18-24 years (n=5), 25-36 years (n=4). (**T**) *Pf*AMA1/*Pf*MSP1 IgM (red) and IgG (blue) reactivity in plasma of 5-10-year-old subjects (n=13), by the MSD assay. Data is expressed as fold change from healthy baseline to convalescence. Statistical analysis: (G-R,S) 2-way ANOVA with Bonferroni’s multiple comparisons test, (T) Wilcoxon matched pairs signed rank test. Lines and error whiskers show means and SDs.

We observed no age-related differences in the proportions of CD21^−^CD27^+^, CD86^+^, CD71^+^ and T-bethi *Pf*^+^ or HA^+^ B cells at the healthy baseline (data not shown). Of note, the response of *Pf*^+^ IgG B cells was similar to the IgM compartment, with an increase in the frequency of CD21^−^CD27^+^, CD86^+^, CD71^+^, Ki67^+^ and T-bethi *Pf*^+^ IgG B cells at convalescence (Fig. 4H, J, N, P and R). Unlike *Pf*^+^ IgM B cells, *Pf*^+^ IgG B cells upregulated HLA-DR (Fig. 4L) in response to malaria. However, in children in particular, the B cell response to febrile malaria was dominated by *Pf*^+^ IgM B cells. This was particularly apparent when the total number of *Pf*^+^ IgM and IgG B cells were enumerated at baseline and convalesence which showed that children under 8 years of age significantly boosted *Pf*^+^ IgM B cells but not *Pf*^+^ IgG B cells (Fig. 4S). With increasing age, *Pf*^+^ IgG B cell numbers at the healthy baseline appear to gradually accumulate, but only exceed IgM B cells in number as individuals reach adulthood (Fig. 4S). Plasma IgM specific for *Pf*AMA1 and *Pf*MSP1 increased 10-fold at convalescence over the healthy baseline (5-10-year-olds, n=13), whereas plasma IgG specific for the same antigens increased 100-fold (Fig 4T).

### Naturally acquired IgM binds to *Pf* antigens and impairs parasite growth *in vitro*

We next investigated whether *Pf*^+^ IgM B cells produce *Pf*-specific IgM antibodies with potentially protective efficacy against blood stage malaria. Direct neutralization of blood stage merozoites, as well as opsonization of parasites for phagocytosis or complement-mediated lysis are possible effector functions of *Pf*-specific IgM. Due to the difficult nature of recombinant expression of IgM, we instead immortalized *Pf*^+^ IgM^+^ B cells (IgD-CD10-CD19^+^ CD20^+^ CD27^+^; see Fig. S3 for sorting strategy) through retroviral expression of Bcl-6 and Bcl-xL (*39*), thereby allowing continuous culture of isolated *Pf*^+^ IgM B cell clones. Culture supernatants containing secreted IgM were tested for reactivity to *Pf*AMA1 and *Pf*MSP1 and several clones with strong binding to *Pf*AMA1 were detected (Fig. 5A). IFAs against *Pf* schizonts confirmed the expected micronemal staining pattern of the *Pf*AMA1-specific IgM monoclonal antibodies (Fig. 5B). Information on VDJ gene usage, CDR3 sequence and number of somatic mutations of cloned hmAbs is displayed in supplementary Table 1. Naturally acquired antibodies have previously been shown to inhibit parasite growth *in vitro* (*40*, *41*) and recently, IgM was found to inhibit merozoite invasion *in vitro*, but only in the presence of complement (*15*). Three IgM clones produced sufficient IgM for GIA (*30*) and affinity-purified anti-*Pf*AMA1 IgM from clone A24 inhibited parasite growth by approximately 60%, compared to 20% inhibition by negative control IgM (Fig. 5C) at a concentration of 1 mg/ml in the absence of complement. To determine whether naturally acquired IgM antibodies from malaria-exposed Malian children and adults have functional activity against *Pf* blood stage parasites, we purified total IgM and IgG from plasma. Tested at equal concentrations (2 mg/ml), both purified total IgM and IgG inhibited parasite growth by an average of 30% compared to 20% inhibition of control IgM obtained from malaria-naïve U.S. donors (Fig. 5D), suggesting that IgM may limit blood stage parasitemia through direct neutralization. The presence of active complement (in the form of non-heat inactivated human serum), did not enhance the inhibitory activity of plasma IgM (Fig. 5D) or IgG (Fig. S4). Moreover, IgM isolated from Malian individuals showed significantly more inhibitory activity than IgM obtained from malaria-naïve U.S. adults (Fig. 5D). Although anti-*Pf*AMA1 and anti-*Pf*MSP1 antibody reactivity increased at convalescence (Fig. 4T), the inhibitory activity of total IgM or IgG was unchanged from healthy baseline to convalescence (Fig. 5D). There was no difference in the inhibitory capacity of total IgM or IgG present in plasma from children and adults (Fig. S4).

**Fig. 5.**
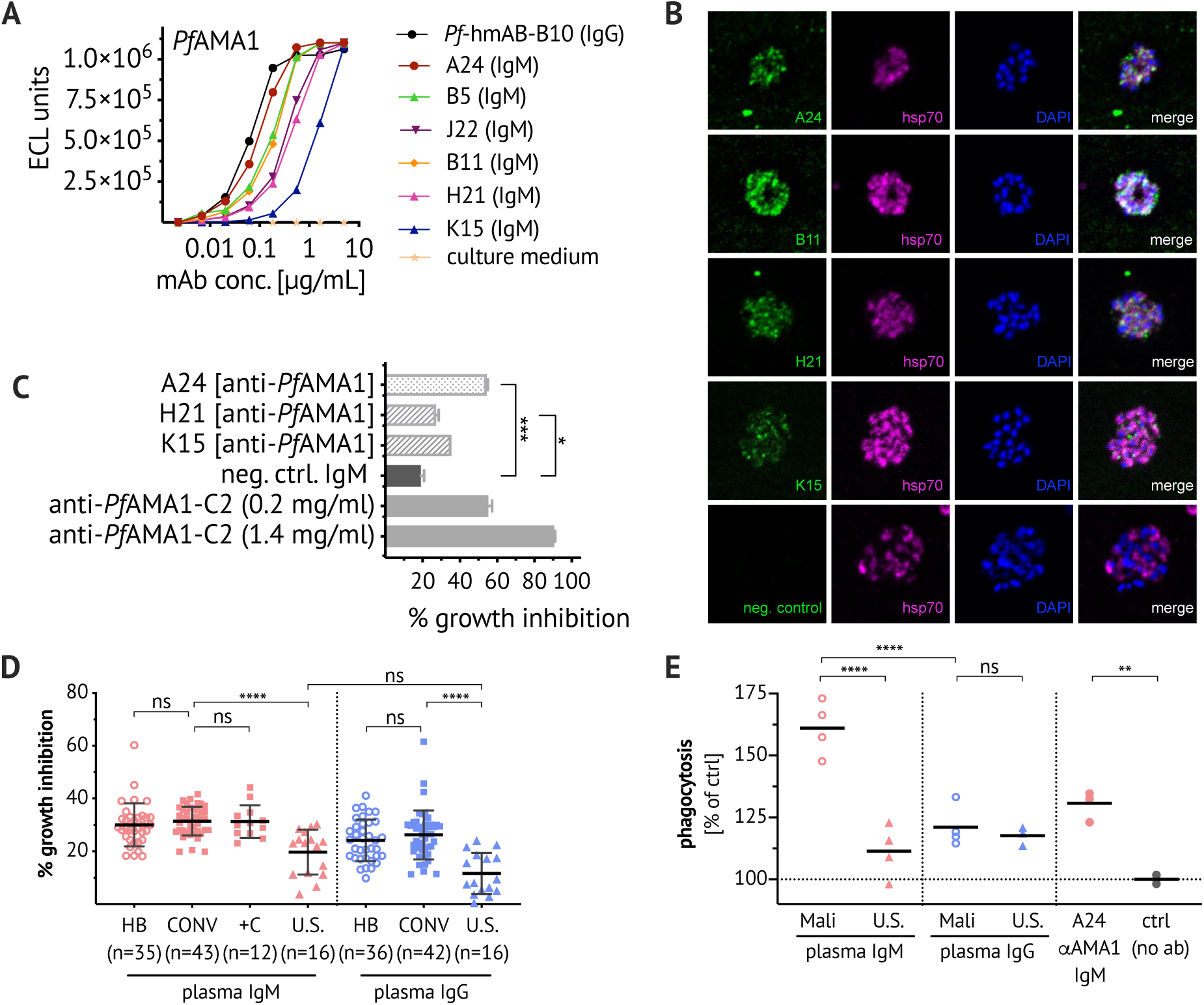
Naturally acquired IgM binds to *Pf* antigens, inhibits growth of *Pf* parasites *in vitro* and enhances phagocytosis of *Pf* merozoite. (**A**) CD19^+^ CD20^+^ IgD-CD10-CD27^+^ IgM^+^ B cells were bulk-sorted for immortalization, followed by single-cell sorting of immortalized *Pf*^+^ B cells for continuous cell culture (see Fig. S3). The MSD assay was used to quantify *Pf*AMA1/*Pf*MSP1-binding of culture supernatant-derived IgM. No strong *Pf*MSP1-binding clones were detected. *Pf*AMA1-binding data is displayed for six IgM clones (A24, B5, J22, B11, H21, K15) and the IgG hmAb clone B10. (**B**) IFA of affinity-purified culture supernatant-derived *Pf*AMA1-specific IgM (clones A24, B11, H21, K15 and negative control clone unrelated specificity). *Pf* 3D7 schizonts were stained with IgM (green, first column), co-stained with anti-hsp70 antibody (purple, second column) and the nuclear stain DAPI (blue, third column). (**C**) Where sufficient quantities were available, *Pf*AMA1-specific IgM (clones A24, H21, K15 and negative control clone) were tested *in vitro* for *Pf* growth inhibitory activity at 1 mg/ml, compared to the positive control polyclonal rabbit anti-*Pf*AMA1-C2 (*31*). Bars and error whiskers show mean and SD. (**D**) Total IgM and IgG were purified from plasma samples obtained from Malian individuals at healthy baseline (HB) and convalescence (CONV) (ages 2-36 years, note that no age-dependent differences were observed, see supplementary Fig. S4) or healthy U.S. adults. Growth inhibitory activity was assessed at 2 mg/ml IgM or IgG. Non-heat inactivated human serum, added at 25% (v/v) as a source of complement (+C), did not enhance the inhibitory effect of IgM. Graph shows data merged from four experiments and data points represent the mean of duplicate wells. Bars and error whiskers show mean and SD. (**E**) pHrodo-Red labelled *Pf* merozoites were pre-incubated with total IgM and IgG (purified from pooled plasma from five U.S. adults, or five malaria-experienced Malian adults), or IgM clone *Pf*-A24. After incubating washed merozoites with THP-1 cells, samples were analyzed by flow cytometry to determine the percentage of pHrodo-Red^+^ THP-1 cells normalized to the percentage pHrodo-Red^+^ THP-1 cells in the absence of antibody. Data from two experiments, each testing samples in duplicates, is shown. Statistical analysis: (C) one-way ANOVA, (D) one-way ANOVA, with Bonferroni’s multiple comparisons test, (E) one-way ANOVA, Tukey’s multiple comparison test.

It has previously been shown that IgM was more effective than IgG at reducing parasite growth in the presence of mononuclear cells (*42*). We tested whether IgM antibodies from malaria-exposed Malians could opsonize parasites for phagocytosis. Total plasma IgM and IgG were isolated from malaria-exposed Malian adults and used to opsonize purified pHrodo-labelled *Pf* merozoites, prior to incubation with THP-1 cells, a human monocytic cell line. As the pHrodo fluorescence intensity increases at lower pH, it provides a stringent measure of phagocytosis into acidic compartments. Plasma IgM from malaria-exposed Malian adults was more effective than plasma IgG at enhancing phagocytosis of merozoites (Fig. 5E), whereas plasma IgM from U.S. adults showed little effect. Together these data suggest that naturally acquired *Pf*-specific IgM contribute to control of parasitemia through direct neutralization and opsonic phagocytosis. Despite these *in vitro* findings, IgM reactivity to *Pf*MSP1 was not associated with reduced risk of febrile malaria in the Mali cohort (Fig. S5).

## Discussion

Present in most vertebrates, IgM is an ancient antibody class that has co-evolved with *Plasmodium* species over millions of years (*18*), yet little is known about the IgM response to malaria, the mechanisms by which IgM may protect, or the underlying biology of *Pf*-specific IgM B cells. Here we analyzed *Pf*AMA1- and *Pf*MSP1-specific B cells (together referred to as “*Pf*^+^“ cells), as well as HA-specific B cells in the peripheral blood of malaria- and influenza-exposed children and adults and found that the majority of non-naïve B cells specific for the blood stage merozoite antigens *Pf*AMA1 or *Pf*MSP1 express IgM, even after many years of repeated malaria infections, in stark contrast to HA-specific B cells that are dominated by IgG B cells. This finding is consistent with a study in mice which showed that *P. chabaudi* infection induces long-lasting somatically mutated IgM MBCs (*19*), as well as recent studies showing that human IgM B cells comprise 30-75% of all CSP-specific B cells in response to both natural *Pf* sporozoite infection and vaccination with the radiation-attenuated PfSPZ vaccine (*43*–*45*).

Although on average, *Pf*-specific IgM BCRs were mutated less than *Pf*-specific IgG BCRs, their level of mutation relative to malaria-naïve controls suggests that they are MBCs. While it has previously been shown that CD27^+^ IgM B cells have somatically mutated variable (V) region genes (*46*), we found it is CD21^−^CD27^−^*Pf*-specific IgM ‘atypical’ B cells that carry V genes with the highest number of mutations. CD21^−^CD27^−^atypical B cells express high levels of T-bet, a transcription factor that has been correlated with IgG3 class switching (*38*), and it may be that T-bet skews B cells towards increased somatic mutation. We further observed that *Pf*-specific IgG BCRs undergo more somatic mutation than HA-specific BCRs, possibly a reflection of the more frequent exposure to *Pf* compared to influenza in the study cohort. HA-specific BCRs cloned from Malian adults had mutation rates similar to HA-specific BCRs cloned from U.S. adults, and both were in the range of previously published mutation data for HA-specific antibodies (*47*). This is interesting, as malaria infections have been found to lead to a disorganized germinal center (GC) architecture in the spleen in rodent malaria models, as well as in *Pf*-infected *Saimiri sciureus* monkeys (*48*). The finding of similar numbers of mutations in influenza-specific antibodies in Malian and U.S. adults indicates that despite potentially frequent alterations of the splenic architecture with repeated malaria episodes, this has no long-term effect on the GC formation in the course of other infections, as measurable through quantification of somatic mutations.

The high prevalence of *Pf*-specific IgM B cells even at the healthy baseline (many months after a febrile malaria episode), despite many years of repeated malaria infections and in absence of asymptomatic parasitemia, indicates incomplete class switching of activated B cells. As discussed previously (*48*), disorganization of the splenic anatomy during the course of a malaria infection may lead to GCs with reduced functionality, which could result in poor *Pf*-specific B cell activation. Additionally, or possibly as a consequence, Th1-polarized PD-1^+^CXCR5^+^CXCR3^+^ T follicular helper (Tfh) cells are preferentially activated during acute malaria (*49*). This subset provided poor B cell help and was less efficient at promoting class switching (*49*, *50*). Furthermore, a recent study showed that TLR9-stimulation leads to a reduced ability of activated B cells to process and present antigen for interaction with antigen-specific helper T cells (*51*). TLR9 ligands, which are abundantly present during malaria infections, such as hemozoin or protein-DNA complexes, may thereby contribute to relativley poor T cell help. Indeed, our data shows that while HLA-DR expression is increased at convalescence from a malaria episode in *Pf*-specific IgM and IgG B cells, this upregulation was statistically significant only in the IgG**^+^** B cell population.

What could be the origin of the *Pf*-specific IgM B cells? Two types of somatically mutated IgM B cells have been described: IgM^+^IgD^+^CD27^+^ marginal zone (MZ)-like B cells, which are thought to be preferentially stimulated in response to blood-borne pathogens present in the splenic marginal zone (*52*); and IgM^+^IgD-CD27^+^ B cells, which are GC-derived and characterized by a higher number of BCR mutations (*53*). We previously found in a small number of malaria-exposed adults that *Pf*-specific IgM^+^CD27^+^ B cells also express IgD, suggesting that *Pf*-specific IgM B cells may be homologous to MZ-like B cells (*19*). Here, our analysis of a larger cohort of malaria-exposed subjects revealed greater complexity of the *Pf*-specific IgM B cell compartment. Specifically, we found that approximately 20% of *Pf*-specific IgM B cells are IgD-, potentially representing IgM MBCs that exited GCs prior to isotype switching. Moreover, just over 40% of *Pf*-specific IgM B cells are IgD^+^CD27^+^ B cells, suggesting that a MZ-like subset may also be activated in the spleen.

We found that *Pf*AMA1- and *Pf*MSP1-specific polyclonal IgG responses in plasma were approxmately 10-fold higher than IgM reponses at the convalescent time point (7 days after treatment of febrile malaria), which may reflect differential kinetics or efficiency of *Pf*^+^ IgM B cells differentiating into antibody-secreting plasmablasts. Due to a low level or lack of surface BCR expression on plasmablasts, our B cell probe method was not directly applicable to the study of isotype distribution in the plasmablast compartment. It is possible that *Pf*^+^ IgM B cells switch to IgG before differentiating into plasmablasts, as was demonstrated for *Plasmodium*-specific MBCs in a malaria rodent model, where *Plasmodium*-specific IgM MBCs efficiently switched to IgG when incubated *in vitro* (*19*). Therefore, human *Pf*^+^ IgM B cells may exhibit similar plasticity and undergo isotype switching prior to differentiation into plasmablasts. In further support of this hypothesis, while the human *Pf*CSP-specific MBC response to vaccination with sporozoites is strongly dominated by IgM, the plasmablast response consisted of comparable proportions of IgA, IgG and IgM (*43*). However, when we analysed 551 *Pf*^+^ IgM and 240 *Pf*^+^ IgG BCR sequences obtained from the same donor, we identified only three clonally related BCRs between the *Pf*^+^ IgM and IgG data sets (Fig. S6), suggesting a limited transition from the IgM to the IgG *Pf*^+^ B cell compartment.

Our data further shows that naturally acquired total IgM can limit parasite growth *in vitro* through direct neutralization of merozoites. Of note, our result differed from a recent study, in which IgM obtained from individuals with a recent malaria infection inhibited merozoite invasion only in presence of active complement (*15*). This difference could be due to the concentration of IgM: while an IgG-depleted fraction was tested for complement-mediated inhibition by IgM at 50 μg/ml (*15*), we tested affinity-purified total IgM at 2 mg/ml, which approximates the IgM concentration in normal plasma (total plasma IgM concentration in Malian individuals was measured and had a median of 2.251 ± 1.4 mg/ml, n=27). Although anti-*Pf*AMA1 and anti-*Pf*MSP1 antibody reactivity increased at convalescence, the inhibitory activity of total IgM or IgG as measured by GIA, was unchanged from healthy baseline to convalescence. The effect of this increase may not be quantifiable by GIA within the total IgM and IgG fractions, or IgM and IgG antibodies targeting parasite antigens other than *Pf*AMA1 and *Pf*MSP1, which may be boosted less by febrile malaria, may be responsible for the growth inhibitory effect of total plasma antibodies.

Opsonization of merozoites by naturally acquired IgM triggered stronger phagocytosis by THP-1 cells *in vitro*, as compared to IgG, suggesting that IgM functions in conjunction with phagocytic cells *in vivo*, through either direct binding of opsonized parasites to FcμR, or complement-mediated opsonization (*54*). In line with this evidence suggesting a protective effect of IgM antibodies are studies that correlated a greater breadth and magnitude of *Pf*-specific IgM, rather than IgG responses with protection against blood stage malaria (*16*, *17*), as well as a recent report showing that high IgM binding to whole merozoites was associated with reduced risk of clinical malaria (*15*). Our study did not reveal an inverse correlation between febrile malaria risk and the IgM titer to *Pf*MSP1, which may be due to our focus on one specific antigen. It would be very interesting to identify the specific merozoite antigens for which IgM reactivity correlates with protection.

As our data shows, *Pf*-specific IgM B cells dominate the *Pf*-specific B cell response at the healthy baseline that precedes the transmission season. These *Pf*-specific IgM B cells expand and express markers of activation and costimulation in response to a febrile malaria episode. We further show that *Pf*-specific IgM B cells produce somatically hypermutated antibodies with antigen-binding characteristics that are comparable to *Pf*-specific IgG antibodies. These data highlight IgM as a significant component of the human immune response to natural malaria infections. While the high prevalence of unswitched *Pf*-specific B cells aids in explaining the inefficient acquisition of a *Pf*-specific IgG response (*7*), this IgM response may also confer advantages in protection from lethal malaria infections through a reduction of blood stage parasitemia: our data suggests that IgM limits parasite growth through direct neutralization and that IgM is superior to IgG in promoting opsonization of merozoites for phagocyte-dependent clearance. Furthermore, having undergone less affinity maturation, IgM might be able to bind a broader range of variant *Pf* antigens. Our data reveals an underappreciated role for IgM in the B cell response to malaria and may stimulate future work toward elucidation of factors that lead to the activation and expansion of this unswitched, but somatically mutated B cell population.

## Methods

### Flow cytometry with *Pf*- and HA-specific B cell probes

The full length ectodomain of apical membrane antigen 1 (*Pf*AMA1) (strain FVO) was expressed in *Pichia pastoris* and the C-terminal 42 kDa region of merozoite surface protein 1 (*Pf*MSP1) (strain FUP) was expressed in *E. coli* as previously described (*25*, *26*). *Pf*AMA1 and *Pf*MSP1 were biotinylated (EZ-link Sulfo-NHS-LC-Biotinylation kit; Thermo Fisher Scientific) using a 1:1 ratio of biotin to protein and once the molar amount of biotinylated protein was measured using a Western blot, proteins were tetramerized with streptavidin-APC (Prozyme) or streptavidin-AlexaFluor488 (Invitrogen) using previously published protocols (*55*). HA probes of two serotypes of influenza virus type A (H1N1 (A/California/04/2009) and H3N2 (A/Texas/50/2012)) were used. HA protein consisting of the extracellular domain of HA C-terminally fused to the trimeric FoldOn of T4 fibritin and the biotinylatable AviTag sequence were expressed and biotinylated as previously described (*27*). HA proteins were tetramerized through sequential addition of streptavidin-BV650 or streptavidin-BV785 (BD biosciences), with HA in excess to streptavidin. For flow cytometric analysis, cryopreserved PBMCs from blood collected at healthy baseline in May or at convalescence from a febrile malaria episode, a week following treatment, were thawed and stained with 100 ng of each B cell probe per sample together with panels using the following labeled mAbs: CD3-BV510 (clone UCHT1), CD4-BV510 (clone SK3), CD8-BV510 (RPA-T8), CD14-BV510 (clone M5E2), CD16-BV510 (clone 3G8), CD56-BV510 (clone HCD56), CD10-APC-Cy7 or CD10-BV510 (clone Hi10a), CD19-ECD (clone J3-119), CD21-PE-Cy7 or CD21-BUV395 (clone B-ly4), CD27^−^BV605 (clone M-T271), IgD-BUV737 (clone IA-2), IgM-BUV395 (clone G20-127), IgG-AlexaFluor700 (clone G18-145), CD71-APC-Cy7 (clone CY1G4), CD86-PE (clone IT2.2), HLA-DR-APC-Cy7 (clone L243), Ki67-PE-Cy7 or KI67-AlexaFluor700 (clone 20Raj1), ICOSL-PE (clone 9F.8A4) and T-bet-PE (clone 4B10). Aqua dead cell stain was added for live/dead discrimination (Thermo Fisher Scientific). Stained samples were run on a LSR II (BD) and data were analyzed using FlowJo (TreeStar). For BCR sequencing, individual live CD3-, CD4-, CD8-, CD14-, CD16-, CD56-, CD19^+^ *Pf*^+^ or HA^+^ B cells were single cell–sorted into 96-well plates using a FACSAria II (BD). Index sorting was used to determine isotype and B cell subsets by IgM, IgG, CD21 and CD27 expression.

### Single-cell BCR amplification, sequence analysis and recombinant hmAb expression

cDNA was directly made from sorted cells using Superscript III Reverse Transcriptase (ThermoFisher) and random hexamers. Immunoglobulin heavy and light chain (kappa and lambda) genes were then PCR amplified separately with two rounds of nested PCR with 50 cycles each, using DreamTaq Mastermix (ThermoFisher), as described previously with some modifications (*22*, *28*). PCR products were Sanger sequenced by Genscript and sequences were analyzed using IMGT/V-QUEST (*33*). Immunoglobulin heavy and light chain sequences were synthesized and cloned by Genscript into IgG1, kappa, or lambda expression vectors. For expression of recombinant hmAbs, Expi293 cells were transfected with plasmids encoding Ig heavy and light chain pairs with Expi-Fectamine (ThermoFisher Scientific) and hmAbs were purified from the culture supernatants using Sepharose Protein A (Pierce).

### *Pf*AMA1/*Pf*MSP1 and HA binding MSD assay

Meso Scale Discovery (MSD) 384 well Streptavidin coated SECTOR Imager 2400 Reader Plates were blocked with 5% MSD Blocker for 1 h, then washed six times with the wash buffer (PBS/0.05% Tween). Streptavidin-plates were then coated with biotinylated *Pf*AMA1 or *Pf*MSP1, H1 or H3 HA protein, as used for flow cytometry, for 1 h and washed. HmAbs and immortalized B cell culture supernatants were diluted in 1% MSD Blocker to 5 μg/ml, serially diluted and added to the coated plates. After a 1 h incubation, plates were washed and incubated with SULFO-TAG conjugated monoclonal anti-human IgG (unknown subclass reactivity) for 1 h. After washing, the plates were read using 1X MSD Read Buffer on the MSD Imager 2400.

### Immunofluorescence assay

Thin blood smears of *Pf* strain 3D7 cultured *in vitro* to the schizont stage as previously described (*56*), were air-dried, fixed with 4% paraformaldehyde/PBS, permeabilized with 0.1% Triton X-100/PBS and blocked with 3% BSA/PBS before incubation with hmAbs at 10 μg/ml and mouse anti-*Pf*hsp70 (clone 2E6, (*57*)) diluted 1:500 in 1% BSA/PBS. Secondary detection was with anti-human AlexaFluor-488 conjugate (A11013, ThermoFisher) and anti-mouse Alexa Fluor 594 conjugate (A11032, ThermoFisher), each diluted 1:1000 in 1% BSA/PBS. Samples were preserved in Prolong Gold mounting medium containing DAPI (Life Technologies), imaged using a LSM880 confocal microscope and images were acquired using Zen software.

### Affinity purification of total plasma IgM and IgG

Columns of 1 ml Protein G sepharose 4 fast flow (GE Healthcare) were prepared and equilibrated with 5 volumes of binding buffer (0.02 M sodium phosphate, pH 7.0) at a linear flow rate of 5 ml/min. 3 ml plasma samples were filtered through glass wool for lipid removal. 3 ml of binding buffer were added, and sample was applied to the protein G sepharose column. Flow-through was collected and used for IgM purification. Column was washed three times with 10 ml binding buffer before IgG was eluted with 5 ml of 0.1 M glycine-HCl, pH 2.7. Eluate was neutralized through addition of 50-100 μl of 1 M Tris-HCl, pH 9.0. For IgM purification, columns of 0.5 ml POROS™ CaptureSelect™ IgM Affinity Matrix (195289005) were prepared and equilibrated with 5 volumes of binding buffer (PBS, pH 7.2-7.4) at a linear flow rate of 5 ml/min. Flow-through from IgG purification was applied and column was washed three times with 10 ml of PBS, pH 7.2-7.4. IgM fraction was eluted with 2.5 ml 0.1 M glycine pH 3.0 and eluate was neutralized with 100-250 μl of 1 M Tris, pH 8.0.

### *Pf in vitro* growth inhibition assay (GIA)

GIAs were conducted with recombinantly expressed hmAbs at final concentrations ranging from 0.0078 to 2 mg/ml or with purified total IgM and IgG at a final concentration of 2 mg/ml using a previously standardized method (*30*). Synchronized *Pf* parasites (3D7 clone) and culture medium were added to 96-well tissue culture plates in duplicate and cultured for ∼40 h. Relative parasitemias were quantitated by biochemical determination of parasite lactate dehydrogenase. Percent inhibition was calculated as 100 - [(A_650_ of test sample - A_650_ of uninfected RBCs)/(A_650_ of control infected RBCs - A_650_ of uninfected RBCs) ×100].

### Transduction of IgM B cells for immortalization

*Pf*-specific IgM B cell clones were generated by expression of Bcl-6 and Bcl-xL in primary B cells, as adapted from a previously described method (*39*). Human codon-optimized Bcl-6 and Bcl-xL were cloned into the pLZRS-IRES-GFP retroviral vector, which was a gift from Lynda Chin (Addgene plasmid #21961; (*58*)). Briefly, live CD10-, CD19^+^, CD20^+^, IgD^−^, CD27^+^, IgM^+^ B cells were isolated from PBMCs on a FACS Aria and activated for 2 days in the presence of recombinant human IL-21 (rhIL-21) (25 ng/mL) and irradiated 3T3-msCD40L cells (*59*) in IMDM + 10% FBS + 1% Penicillin-Streptomycin-Glutamine (Gibco) (I10). Activated B cells were centrifuged with viral supernatant containing 4 μg/ml polybrene (1200*xg*, 32C, 1 h) and subsequently cultured in I10 with 3T3-msCD40L cells and rhIL-21 at 37°C. After 3 days, transduced clones were stained with *Pf* probes, and the *Pf*^+^ GFP^+^ B cells were sorted into 384-well plates containing I10, rhIL-21 and 3T3-msCD40L cells. Following long-term culture, supernatants were screened by ELISA for Ig binding to *Pf*AMA1 and *Pf*MSP1. To expand clones for IgM production, cells were cultured in serum-free hybridoma media (ThermoFisher) in the presence of IL-21 and irradiated msCD40L feeder cells and IgM was purified from supernatant using POROS™ CaptureSelect™ IgM Affinity Matrix (195289005).

### Phagocytosis of *Pf* merozoites by THP-1 cells

*Pf* merozoites were harvested from supernatant of *Pf* strain 3D7, cultured to the schizont stage as previously described (*56*). The parasite culture was centrifuged at 400*xg* for 10 min to pellet schizonts and uninfected RBCs. The supernatant was collected and spun at 3200*xg* for 10 min to collect merozoites in the pellet, which were cryopreserved in 2% DMSO in FCS. Purified *Pf* merozoites were stained with pHrodo ™ iFL Red STP amine-reactive ester (ThermoFisher) at 8 μM in RPMI and incubated to allow for opsonization with 2 mg/ml plasma IgM or IgG antibodies for 1 h at RT. After a washing step, phagocytosis was performed by incubation of merozoites with THP-1 cells (ATCC) at a ratio of 1 to 10 for 2 h at 37°C h in RPMI with 5% albumax, in a 5% CO2 humidified incubator. A control performed at 4°C, as well as a sample with no merozoites were included. Samples were analyzed by flow cytometry, and phagocytosis was determined as the percentage of THP-1 cells that were pHrodo-Red^+^ and normalized to percentage of phagocytosis in presence of no antibody.

### Cross-sectional quantification of plasma IgM binding to *Pf*MSP1

Streptavidin-coated beads with different levels of FITC-channel fluorescence were incubated with 10 μg/ml of biotinylated *Pf*MSP1 (*60*) or biotinylated rat CD4 as a negative control antigen for 30 minutes at RT. The beads were washed, blocked with a further 10 μg/ml of biotinylated rat CD4, and mixed. They were then incubated with 1/30 of plasma from the Malian cohort, followed by incubation with 1/150 BUV395 anti-human IgM (BD) and analysis by flow cytometry. The amount of IgM binding was calculated as the signal ratio of *Pf*MSP1 to the negative control antigen (after correction through subtraction of the fluorescence of the beads with no serum added).

## Acknowledgements

We thank the residents of Kalifabougou, Mali, for participating in this study. *Pf*AMA1 and *Pf*MSP1 proteins were kindly provided by David Narum at the Laboratory of Malaria Immunology and Vaccinology/NIAID. We are grateful to Gavin Wright and Nicole Muller-Sienerth at the Wellcome Sanger Institute for sharing recombinant *Pf*MSP1. We thank Justin Taylor (Fred Hutchinson Cancer Research Center) for helpful advice on B cell probe development and Photini Sinnis (Johns Hopkins Bloomberg School of Public Health) for the generous gift of the hsp70 antibody and Tongqing Zhou from Peter Kwong’s lab (VRC, NIAID) for providing IL-21 used in the retroviral expression experiments. This work was supported by the Division of Intramural Research of the National Institute of Allergy and Infectious Diseases, National Institutes of Health.

## Supplementary Materials

**Table S1.** V, D and J gene information, CDR3 sequence and the number of amino acid (aa) changes of cloned *Pf-*and HA-specific hmAbs and *Pf-*specific immortalized IgM B cell clones.

**Fig. S1.** Plasma reactivity to influenza HA in children and adults in Kalifabougou, Mali.

**Fig. S2.** Comparable subset distribution within switched and unswitched *Pf*-specific IgD-B cell populations and high numbers of BCR mutations in atypical *Pf*-specific IgM B cells.

**Fig. S3.** Sorting strategies for BCR sequencing and immortalization of *Pf*-specific IgM B cells.

**Fig. S4.** IgM inhibition of *Pf* parasite growth *in vitro* shows no age-dependent differences.

**Fig. S5.** Plasma IgM reactivity to *Pf*MSP1 in children and adults in Kalifabougou, Mali.

**Fig. S6.** Circos plot showing clonal relationships shared between IgM and IgG *Pf*-specific BCR sequences obtained from one donor.

## Notes

### Competing Interest Statement

The authors have declared no competing interest.

