## Supplemental Material for "*Plasmodium falciparum*-specific IgM B cells dominate in children, expand with malaria and produce parasite inhibitory IgM"

| *Pf* mAb (IgG) | memory subset | SHM (aa changes) | heavy chain V-gene and allele | heavy chain D-gene and allele | heavy chain J-GENE and allele | heavy chain CDR3 | light chain V-GENE and allele | light chain J-GENE and allele | light chain CDR3 |
|---|---|---|---|---|---|---|---|---|---|
| ***Pf*-C3** | **activated** | 2 | IGHV3-30*02 F | IGHD3-16*01 F | IGHJ6*04 F | GRSGAKWASEPMDV | IGLV1-47*01 F | IGLJ2*01 F or IGLJ3*01 F | SVWDDNLNGLV |
| ***Pf*-C11** | **classical** | 1 | IGHV3-30*02 F | IGHD3-9*01 F | IGHJ6*04 F | AKVGWTWVSEPADV | IGLV1-47*01 F | IGLJ2*01 F or IGLJ3*01 F | AVWDGDLSGVI |
| ***Pf*-D6** | **immature/naïve** | 5 | IGHV4-4*07 F | IGHD3-22*01 F | IGHJ4*02 F | AREIYYHDSTGSLYYFDY | IGKV3-15*01 F | IGKJ1*01 F | LQFNDWPPT |
| ***Pf*-B10** | **atypical** | 3 | IGHV4-61*02 F | IGHD2-15*01 F | IGHJ3*01 F | AKESGCDGGICYGPLYV | IGKV4-1*01 F | IGKJ1*01 F | QQYYSSPRT |

| *Pf* immortalized clone (IgM) | memory subset | SHM (aa changes) | heavy chain V-gene and allele | heavy chain D-gene and allele | heavy chain J-GENE and allele | heavy chain CDR3 | light chain V-GENE and allele | light chain J-GENE and allele | light chain CDR3 |
|---|---|---|---|---|---|---|---|---|---|
| ***Pf*-A24** | **classical** | 30 | IGHV4-59*02 F | IGHD7-27*01 F | IGHJ5*02 F | SRAWDR | IGKV3-20*01 F | IGKJ5*01 F | QQRGGSGVT |
| ***Pf*-B5** | **classical** | 30 | IGHV4-59*02 F | IGHD7-27*01 F | IGHJ5*02 F | SRTWDR | IGKV3-20*01 F | IGKJ5*01 F | QQRGGSGVT |
| ***Pf*-B11** | **classical** | 29 | IGHV4-59*02 F | IGHD7-27*01 F | IGHJ5*02 F | SRTWDR | IGKV3-20*01 F | IGKJ5*01 F | QQRGGSGVT |
| ***Pf*-J22** | **unkown** | 16 | IGHV3-66*02 F | IGHD4-23*01 | IGHJ4*02 F | ASGPLSILRGPD | IGKV3-20*01 F | IGKJ1*01 F | HQYGKSPWT |
| ***Pf*-H21** | **unkown** | 17 | IGHV3-66*02 F | IGHD4-23*01 | IGHJ4*02 F | ASGPLSILRGPD | IGKV3-20*01 F | IGKJ1*01 F | HQYGKSPWT |
| ***Pf*-K15** | **unkown** | 7 | IGHV5-51*05 F | IGHD5-24*01 | IGHJ1*01 F | ARRKASGHNYSFQH | IGLV3-25*03 F | IGLJ2*01 F | SINRQQWYLCG |

| HA mAb (IgG) | memory subset | SHM (aa changes) | heavy chain V-gene and allele | heavy chain D-gene and allele | heavy chain J-GENE and allele | heavy chain CDR3 | light chain V-GENE and allele | light chain J-GENE and allele | light chain CDR3 |
|---|---|---|---|---|---|---|---|---|---|
| **HA-G9** | **classical** | 8 | IGHV3-21*01 F | IGHD3-3*01 F | IGHJ6*03 F | ARDSGIKSNFDFWTGFHYFYYMDV | IGLV7-43*01 F | IGLJ2*01 F | LLYYGGAQV |
| **HA-B11** | **classical** | 19 | IGHV4-61*02 F | IGHD3-9*01 F | IGHJ3*02 F | ARRDYDILTGGGNDAFDI | IGLV1-44*01 F | IGLJ2*01 F | AVWDDSLNGVV |
| **HA-D5** | **immature/naïve** | 4 | IGHV1-18*01 F | IGHD3-22*01 F | IGHJ6*02 F | ARSGYYDGSNYYSYDYYGLDV | IGKV3-15*01 F | IGKJ1*01 F | QQYNDWLRT |
| **HA-D9** | **atypical** | 8 | IGHV1-69*01 F | IGHD2-15*01 F | IGHJ3*02 F | ARREGYCSGGSCYSSFHI | IGLV1-44*01 F | IGLJ1*01 F | AAWDGSLN |

**Table S1.** V, D and J gene information, CDR3 sequence and the number of amino acid (aa) changes of cloned *Pf*- and HA-specific hmAbs and *Pf*-specific immortalized IgM B cell clones.

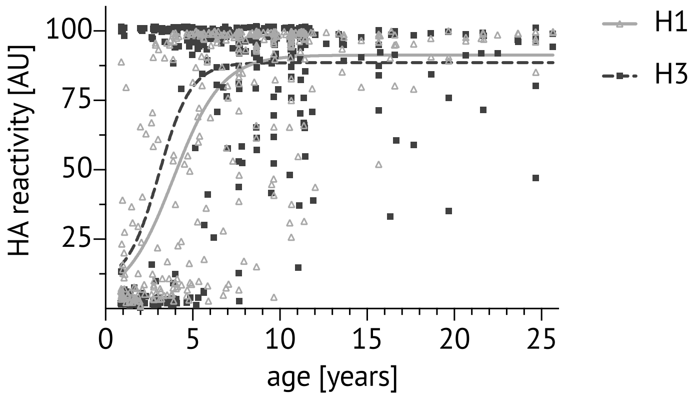


**Fig. S1. Plasma reactivity to influenza HA in children and adults in Kalifabougou, Mali.** Cross-sectional analysis in May 2011 of plasma binding to HA probes of H1N1 and H3N2 subtypes using the MSD assay. Dots show the individual sample antibody level in arbitrary units (AU), calculated as a percentage of the positive control hmAb CR9114. Lines denote the non-linear fit of reactivity to H1 (solid light grey line) and H3 (dotted line).

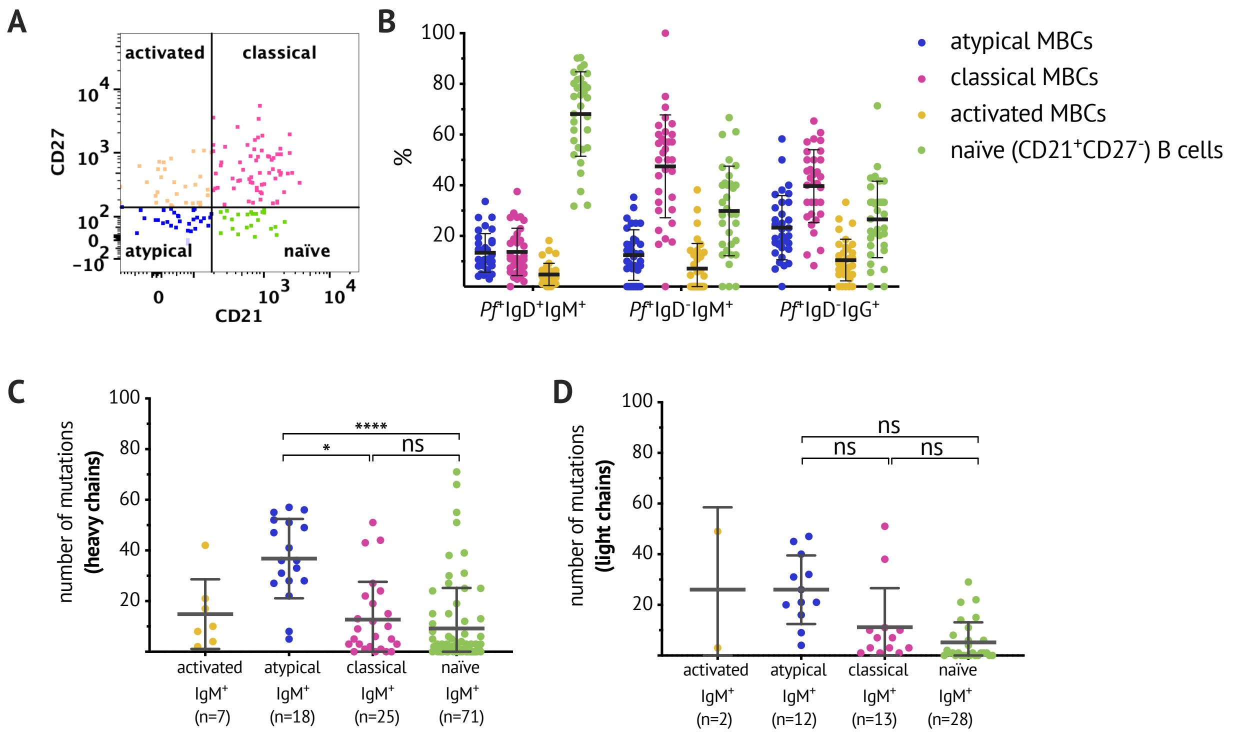


**Fig. S2. Comparable subset distribution within switched and unswitched *Pf*-specific IgD$^-$ B cell populations and high numbers of BCR mutations in atypical *Pf*-specific IgM B cells.** (**A**) Flow cytometry gating strategy to identify B cell subpopulations. (**B**) Distribution of naïve B cells (CD21$^+$CD27$^-$), classical (CD21$^+$CD27$^+$), atypical (CD21$^-$CD27$^-$) and activated MBCs (CD21$^-$CD27$^+$) as a percentage of *Pf*$^+$IgD$^+$IgM$^+$ B cells, *Pf*$^+$IgD$^-$IgM$^+$ B cells or *Pf*$^+$IgD$^-$IgG$^+$ B cells. Each dot indicates an individual; lines and whiskers represent means and SDs. (**C-D**) Number of mutations in the variable region of the heavy chain (**C**) and light chain (**D**) of individual *Pf*$^+$ IgM$^+$ B cells. Data combined from five healthy adult Malian donors. Each dot indicates a single cell; lines and whiskers represent means and SDs. Statistical analysis: (C-D) One-way ANOVA with Holm-Sidak's multiple comparisons test.

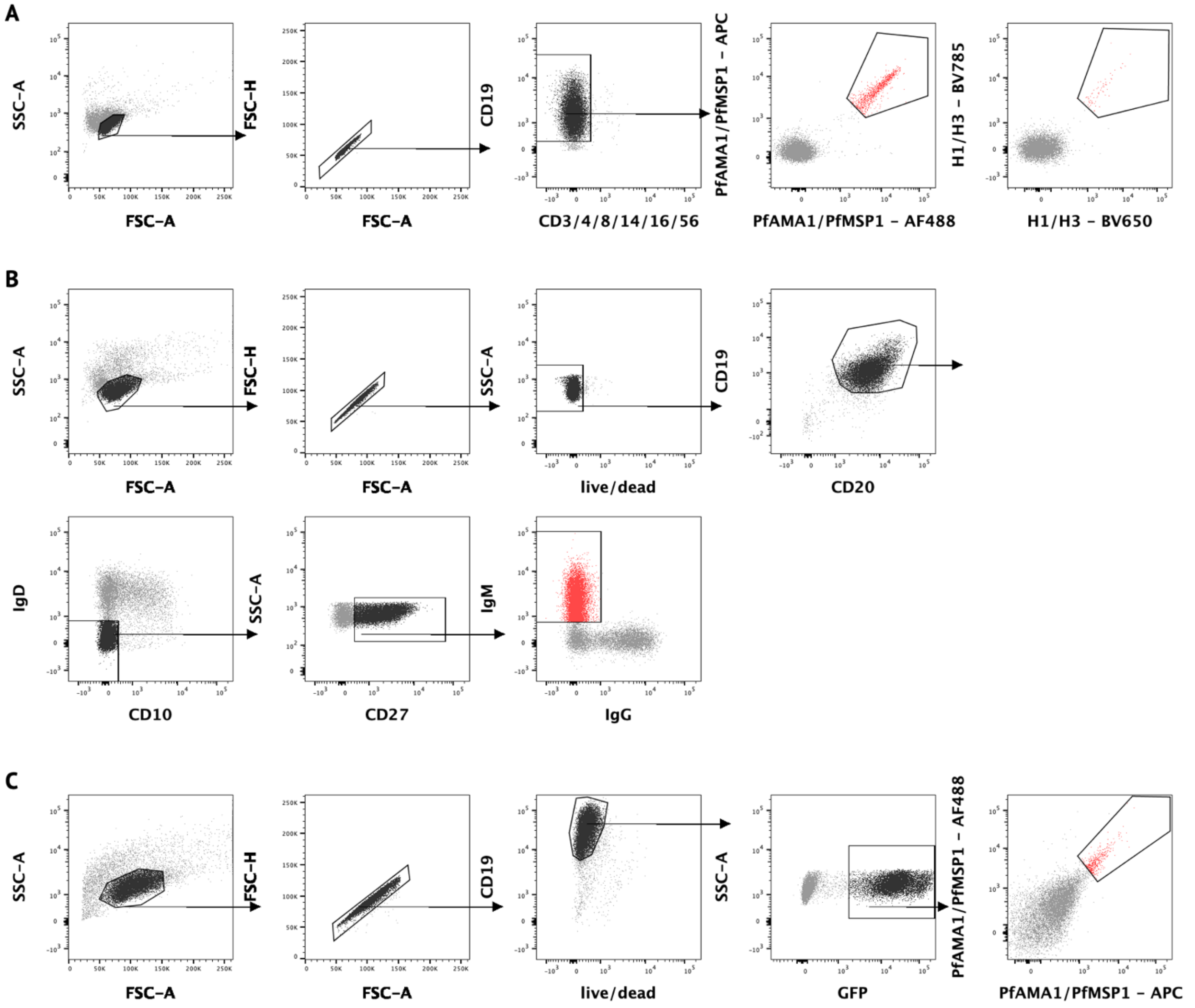


**Fig. S3. Sorting strategies for BCR sequencing and immortalization of *Pf*-specific IgM B cells.** Representative flow cytometry plots of Malian PBMCs after B cell enrichment, showing gating on (**A**) live CD3$^-$, CD4$^-$, CD8$^-$, CD14$^-$, CD16$^-$, CD56$^-$, CD19$^+$ *Pf*$^+$ or HA$^+$ B cells that were single cell–sorted for BCR sequencing, and (**B**) live CD19$^+$ CD20$^+$ IgD$^-$ CD10$^-$ CD27$^+$ IgM$^+$ B cells that were bulk sorted for transduction with the pLZRS-IRES-GFP retroviral vector expressing Bcl-6 and Bcl-xL (a gift from Lynda Chin, Addgene plasmid #21961; (*58*)). (**C**) Five days after transduction, B cells were stained with *Pf* probes. Representative flow cytometry plots of transduced B cells showing gating on live CD19$^+$ GFP$^+$ B cells expressing GFP. *Pf*AMA1 or *Pf*MSP1 probe-binding cells were single-cell sorted into 384-well plates for continuous *in vitro* culture and BCR sequencing.

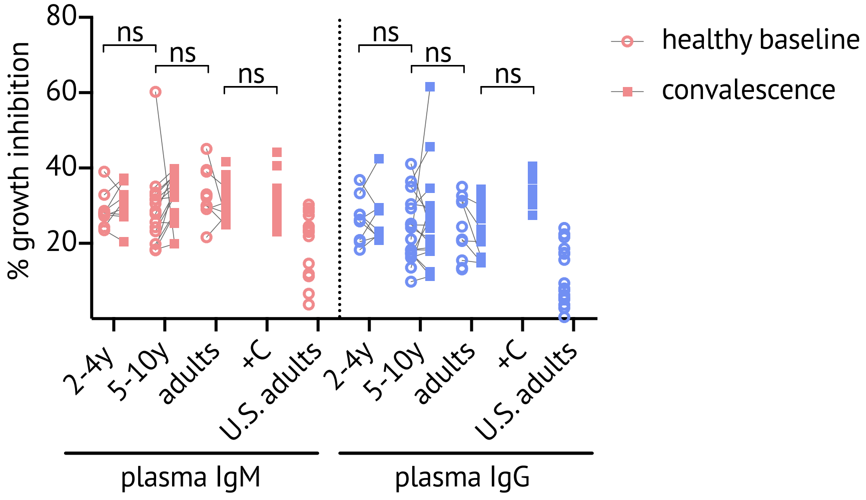


**Fig. S4. IgM inhibition of *Pf* parasite growth *in vitro* shows no age-dependent differences.** Total IgM and IgG were purified from plasma samples obtained from Malian children and adults (ages 2-4 years (n=9), 5-10 (n=16), adults (n=10) at healthy baseline (circles) and convalescence (squares) and from U.S. adults (n=16). Growth inhibitory activity was assessed at 2 mg/ml for both IgM and IgG. Non-heat inactivated human serum was added at 25% (v/v) as a source of complement (+C) and did not enhance growth inhibitory effect of purified plasma IgM or IgG (n=12). Data points represent the mean of duplicate wells. Statistical analysis: paired (and unpaired) comparisons were made with a nested design, linear mixed model ANOVA and Tukey-adjusted post hoc tests.

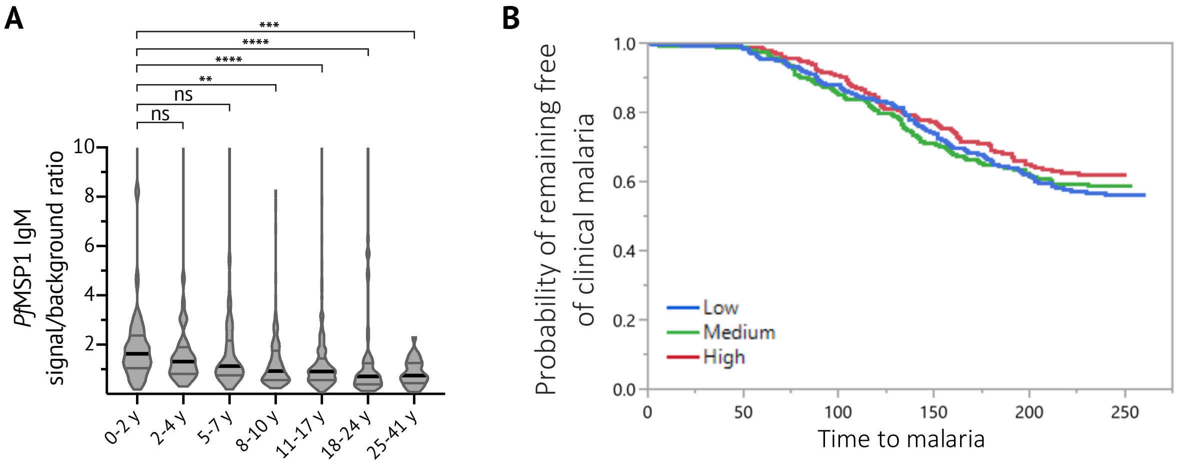


**Fig. S5. Plasma IgM reactivity to *Pf*MSP1 in children and adults in Kalifabougou, Mali.** (**A**) Cross-sectional analysis of IgM binding to *Pf*MSP1 (*60*) at healthy baseline is shown for individuals aged 0-2 years (n=66), 2-4 years (n=37), 5-7 years (n=88), 8-10 years (n=103), 11-17 years (n=378), 18-24 years (n=58) and 25-41 years (n=28). IgM binding was quantified as the signal ratio of *Pf*MSP1 to the negative control antigen (CD4). Data is displayed in violin plots with black line indicating the median and grey lines highlighting the interquartile ranges. (**B**) Individuals were stratified into three equal-sized tertile groups according to low, medium or high *Pf*MSP1 IgM signal ratio. Statistical analysis: (A) Kruskal-Wallis, Dunn's multiple comparison test, (B) Tertile groups were compared for time to the first febrile malaria episode during the ensuing transmission season by Kaplan-Meier survival analysis (Log-Rank $p = 0.4829$, Wilcoxon $p = 0.4446$). A Cox Proportional Hazards regression looking at subject age (Wald $p < 0.0001$), Log10 *Pf*MSP1 IgM titer (Wald $p = 0.9820$) and their interaction (Wald $p = 0.1680$) did not show any significant effect of *Pf*MSP1 IgM reactivity.

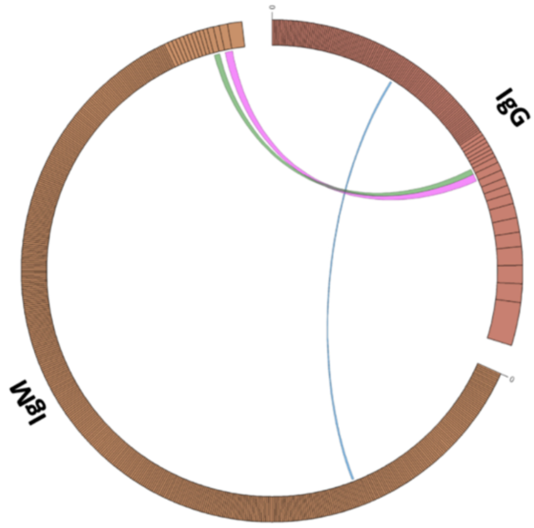


**Fig. S6. Circos plot showing clonal relationships shared between IgM and IgG *Pf*-specific BCR sequences obtained from one donor.** 551 *Pf*$^{+}$ IgM and 240 *Pf*$^{+}$ IgG BCR sequences were obtained through single cell and bulk sequencing of flow cytometry sorted *Pf*$^{+}$ IgM and IgG B cells obtained from one Malian malaria-exposed donor. From this single donor PBMCs were pooled from healthy baseline and convalescence timepoints from 2015-2018. Circos plots show all sequences for IgG and IgM cells in separate segments, divided into clones (based on same V, same J, same CDR3 length, and 85% similarity of the CDR3 nucleotide sequence), and ranked from the largest clones in the clockwise-most position. Lines on the interior of the plots connect matching clones found in both isotypes. Three clones were found to be related between the IgM and IgG data sets, amounting to 1.9% of the IgG clones with a matching IgM clone.
